# Structural insight into the transcription activation mechanism of the phage Mor/C family activators

**DOI:** 10.1101/2025.05.02.651988

**Authors:** Jing Shi, Zonghang Ye, Yirong Huang, Liqiao Xu, Simin Xu, Lihong Xie, Wei Chen, Lu Wang, Zhenzhen Feng, Qian Song, Shuang Wang, Yu Feng, Wei Lin

## Abstract

Bacteriophage Mu, a temperate phage that infects *E*. *coli* K-12 and other enteric bacteria, precisely controls its replication cycle through hijacking host RNA polymerase (RNAP) by the middle operon regulator Mor and the late gene transcription activator C. Though a dimeric arrangement and significant conformational changes are proposed for the distinct Mor/C family activators, the underlying transcription activation mechanism remains unclear. In this study, we present two cryo-EM structures of the transcription activation complex (Mor-TAC and C-TAC) with phage Mu middle and late gene promoters, respectively. Remarkably, the Mor/C activators bind to promoter DNA as a centrosymmetric tetramer rather than as the proposed dimer, concurrently stabilizing by the N-terminal dimerization domains and C-termini. The C-terminal DNA binding domains and two anti-β-strands simultaneously interact with two adjacent DNA major grooves. The activators also engage a variety of interactions with the conserved domains (αCTD, σ^70^R4, and β FTH) of RNAP, providing evidences for a recruitment mechanism. In addition, single-molecule FRET assays show that C significantly enhances RPitc formation, suggesting a different multi-step activation mechanism for C. Collectively, these findings reveal the unique transcription activation mechanism of tetrameric Mor/C family activators, unraveling a novel mode of phage hijacking and bacterial transcription regulation.

## INTRODUCTION

Though encompassing simple regulatory systems, the ubiquitously distributed phages possess the capability to resist bacterial adversities and govern phage replication cycles (1). They primarily achieve this by encoding various transcription factors that hijack bacterial transcriptional machineries via intricate protein-DNA and protein-protein interactions during the transcription initiation process (2,3). In the model bacteria *Escherichia coli* (*E*. *coli*), the multi-subunit RNA polymerase holoenzyme (RNAP, α2ββ’ωσ) is initially assembled by the core enzyme and the principal promoter specific factor σ (σ^70^) (4,5). Accompanied by interacting with the β flap and the clamp helices of β’ subunit (β’ CH), the conserved domains R4 (σ^70^R4) and R2 (σ^70^R2) of σ^70^ specifically recognize promoter DNA consensus -35 and -10 elements, respectively (6). This drives DNA unwinding and isomerizes an RNAP-promoter closed complex (RPc) into an RNAP-promoter open complex (RPo) (7–9). With the addition of NTPs, RNAP attempts to synthesize nascent RNA short than 12 nucleotides, result in an RNAP-promoter initial transcribing complex (RPitc). As RNA length reaches 13–15 nucleotides, σ^70^ disengages from interactions with the promoter DNA and RNAP—a process known as promoter escape (or promoter clearance)—thereby transitioning abortive transcription initiation into the subsequent transcription elongation stage (10–12). For weak promoters that consist of suboptimal promoter elements, various transcription activators join in to compensate for instability between the RNAP and promoter DNA, accumulating formation of a transcription activation complex (TAC) competent for transcription initiation (13–18).

Bacteriophage Mu is a temperate phage that infects *E*. *coli* K-12 and other enteric bacteria. To ensure successful infection, Mu has evolved a complex and elaborate regulatory network to control transient expression of distinct phase genes (early, middle, and late genes) and to switch on the lytic cycle (19–22). Notably, phage Mu lacks self-encoded RNA polymerase and primarily relies on host RNAP and phage-encoded activators to control the weaker promoters of middle genes and late genes, making Mu a valuable model system for studying phage hijacking strategies and bacterial transcriptional regulation. The middle operon regulator (Mor) activates transcription of the middle gene promoter (P*m*) which controls the expression of the late transcription activator C (22–25). While C activates transcription of four late promoters (*P_mom_*, *Plys*, *P_I_*, *P_P_*) that regulate phage DNA modification, morphogenesis, and cell lysis (20,21,26–29). Both structural and bioinformatics analyses reveal that Mor and C exhibit significant structural similarities, including an acidic N-terminal dimerization domain (DD), a conserved 12-amino acid β-strand linker, and a basic C-terminal DNA binding domain (DBD) (23,30–35)(**Fig. S1**). Interestingly, the Mor/C proteins, along with the Mu-like phage proteins, form a structurally distinct Mor/C family, which has been extensively investigated for decades. Yet, the molecular mechanisms underlying transcription activation by these activators remain elusive.

Biochemical assays have shown that both Mor and C proteins form dimers in solution, which may enable dimeric binding to the imperfect dyad-symmetry elements in middle and late genes of phage Mu, further recruiting RNAP to promoter DNA (23,30,35,36). While the crystal structure and mutational analyses of Mor have highlighted the significance of hydrophobic interactions in stabilizing the dimerization domain (DD) and the essential role of the helix-turn-helix (HTH) motif in engaging with two adjacent major grooves of DNA, substantial conformational changes have also been proposed to address the considerable discrepancies observed between the docked model of Mor-DNA and previous chemical cross-linking experiments (30,32,36,37). The DNA bending angle induced by Mor binding and the adjacent linker, has been suggested to contribute to this conformation change (30,32). Furthermore, biochemical and genetic assays have indicated that the C-terminal domain of RNAP α subunit (αCTD) and the C-terminal R4 domain of σ^70^ (σ^70^R4) are required for Mor-dependent transcription of middle genes (25,38–42). Despite the high structural conservation of Mor, it has been found that C is critical for the creation of productive TAC, inducing allosteric transitions, and enhancing efficient promoter escape via a multistep transcription activation mechanism on phage late gene promoters (30,31,43,44). Nonetheless, the lack of structural information regarding the binary Mor/C-DNA complex or the ternary Mor/C-DNA-RNAP complex leaves long-standing questions. Additionally, the role of the short N-terminus (26 amino acids) and C-terminus (9 amino acids) unresolved in the crystal structure remains an intriguing query that warrants further exploration.

In the present work, we report the cryo-EM structures of phage Mu Mor-dependent transcriptional activation complex (Mor-TAC, 4.12 Å) and C-dependent transcriptional activation complex (C-TAC, 3.14 Å), which are assembled on phage Mu middle and late gene promoters, respectively. Strikingly, in both Mor-TAC and C-TAC, the Mor and C proteins bind to promoter DNA as a centrosymmetric tetramer, with the N-terminal DDs and C-termini facilitating this unique architecture. The C-terminal DNA binding domains and two anti-β-strands concurrently interact with the two adjacent DNA major grooves upstream of promoter -35 element, while two flanking helices α2 make contacts with the bilateral DNA minor grooves. Furthermore, the stability of TACs is further stabilized through diverse interactions between the Mor/C activators and the conserved domains (αCTD, σ^70^R4, and β FTH) of RNAP, providing structural evidences for the recruitment mechanisms for Mor and C. Consistently, mutagenesis and single-molecule FRET assays proved C significantly enhances RPitc formation, thus supporting a different multi-step transcription activation mechanism for C. Altogether, these data offer molecular insights into the unique transcription activation mechanisms of tetrameric Mor/C family activators, unraveling a novel mode of phage hijacking and bacterial transcription regulation.

## MATERIAL AND METHODS

### Preparation of plasmids and DNA

Plasmids of pET28a-*mor* carrying N-terminal 6*His tagged phage Mu middle gene activator Mor encoding gene *mor* and pET28a-*c* carrying N-terminal 6*His tagged phage Mu late gene activator C encoding gene *c*, both under the control of T7 promoter, were synthesized by Sangon Biotech, Inc. The P*m* DNA fragment harboring the phage Mu middle gene promoter Pm (-87 to +30, including Mor binding box) fused with a following RNA aptamer mango III encoding sequence *mango* (*45,46*) was amplified by de novo PCR by using primers of *Pm_F*, *Pm_F1*, *Pm_R1*, and *mango_R* (**Table S1**). Similarly, the P*mom* DNA fragment harboring the phage Mu late gene promoter Pm (-81 to +30, including C binding box) fused with a following *mango* sequence was prepared by using primers of *mom_F*, *mom _F1*, *mom_R1*, and *mango_R* (**Table S2**). Plasmids of pGEX-4T-Mor and pGEX-4T-C were constructed by amplifying target genes and pGEX-4T-1 vector fragment using the corresponding primers (**Tables S1** and **S2**) and through homologous recombination methods according to the manual instructions (Vazyme, Inc). All of the other primers used for constructing mutants of Mor and C are listed in **Tables S1** and **S2**.

### Purification of Phage Mu Mor and C

*E*. *coli* phage Mu late gene activator C was transformed with the plasmid pET28a-*c* or pET28a-*c* derivatives. The identified positive transformants were firstly inoculated at 37 °C until OD600 reached 0.8, and induced with 0.5 mM IPTG at 18 °C for 16 h. The cells were then harvested, resuspended, and lysed in buffer A (20 mM Tris-HCl, pH 8.0, 200 mM NaCl, 5% glycerol). The supernatant centrifuged at 12800 g for 30 min was loaded onto a 3 ml Ni-NTA agarose column (Qiagen, Inc.) equilibrated with buffer A. The column was sequentially washed with 15 ml buffer A containing 25 mM, 40 mM imidazole and eluted with 15ml buffer A containing 200 mM imidazole. The targeted fractions containing Mor were verified by SDS-PAGE and concentrated for further analysis. The derivatives of C were purified and stored as the wild type (WT) proteins. By analogy, the GST-C was loaded and eluted from the glutathione agarose resin column and further purified to high homogeneity by a HiLoad 16/600 Superdex 75 column (GE Healthcare, Inc.) in buffer B (20 mM Tris-HCl, pH 8.0, 75 mM NaCl, 5 mM MgCl2, 1 mM DTT).

Likewise, *E*. *coli* phage Mu middle gene activator Mor was transformed with the plasmid pET28a-*mor* or pET28a-*mor* derivatives. Then the Mor proteins and its derivatives were prepared to high purity through similar affinity chromatography columns and gel filtration chromatography column as described above for C, which allows for subsequent experimental analysis.

### Purification of *E. coli* RNAP

*E. coli* RNAP was prepared from *E. coli* strain BL21(DE3) (Invitrogen, Inc.) transformed with plasmids of pGEMD (47) and pIA900 (48) by following the methods as described previously (49,50) and indicated in the present work. Single positive colonies were used to incubate, amplify, and induce protein expression by addition of 0.5 mM IPTG at 18 °C for 16 h. After harvest by centrifugation (6000 g; 15 min at 4 °C), the cell pellets were resuspended and lysed in 20 ml lysis buffer A according to per liter of LB medium. Then the supernatant centrifuged was sequentially processed through Polyethylenimine (PEI) precipitation (w/v, 0.7%), washing with buffer C (20 mM Tris–HCl pH 7.9 and 5% glycerol) containing 500 mM NaCl for three times, extraction with buffer C containing 1000 mM NaCl, ammonium sulfate precipitation (w/v, 30%), resuspending with buffer C containing 500 mM NaCl, and loading onto a pre-equilibrated Ni-NTA agarose (Qiagen, Inc.) column. After washing with buffer C containing 10 mM imidazole, the column eluted with buffer C containing 200 mM imidazole. Subsequently, the eluate was further purified through a Mono Q 10/100 GL column (GE Healthcare, Inc.) with a 160 ml linear gradient of 300-500 mM NaCl in buffer D (20 mM Tris-HCl, pH 7.9, 5% glycerol, 1 mM EDTA and 1 mM DTT). Target fractions carrying *E. coli* RNAP were applied and purified to high homogeneity by a HiLoad 16/600 Superdex 200 column (GE Healthcare, Inc.) in buffer B. Fractions were pooled and concentrated to 7.6 mg/ml.

### Assembly of phage Mu Mor–TAC and C-TAC

The template strand DNA (T) and nontemplate strand DNA (NT) of P*m s*caffold and P*mom s*caffold were synthesized by Sangon Biotech, Inc, and annealed at 1:1 ratio in 10 mM Tris– HCl, pH 7.9, 0.2 M NaCl. Then, the assembly of the C-TAC or Mor-TAC was initiated by incubating *E. coli* RNAP, P*mom s*caffold (or P*m s*caffold), and phage Mu C protein (or Mor protein) in a molar ratio of 1: 1.1: 8 at 37 °C for 10 min, and then at 4 °C overnight. After centrifugation, the resultant supernatant was loaded onto a HiLoad 16/600 Superdex 200 column (GE Healthcare, Inc.) equilibrated in buffer B, and the column was eluted with 120 mL of the same buffer. Following verification via SDS-PAGE, the target fractions containing each component of the assembled C-TAC or Mor-TAC were concentrated to 20.6 mg/ml (24.5 mg/ml) by using Amicon Ultra centrifugal filters (10 kDa MWCO, Merck Millipore, Inc.). The Oligonucleotides synthesized for the preparation of the DNA scaffolds are detailed in **Tables S1** and **S2**.

### Cryo-EM grid preparation

Initially, the Quantifoil grids (R1.2/1.3 Cu 300 mesh holey carbon grids; Quantifoil, Inc.) underwent glow_discharge for 120 seconds at a current of at 15 mA. Following incubation with 6 mM CHAPSO (Hampton Research Inc.) for 1 min at 25 ℃, 3 μL of the phage Mu C-TAC (or Mor-TAC) sample was loaded onto the grids. After blotting with Vitrobot Mark IV (FEI), the grids were immediately plunge-frozen in liquid ethane with 95 % chamber humidity at 10 ℃. Finally, grids exhibiting a moderate density and uniform distribution of single particles were selected for extensive cryo-EM data collection.

### Cryo-EM data collection and processing

Cryo-EM data for C or Mor-dependent transcription activation complex was acquired using a consistent set of parameters on a 300 kV Titan Krios (FEI, Inc.) equipped with a K3 Summit direct electron detector. The data were processed sequentially with the appropriate cryo-EM data analysis software including CryoSPARC v4.2 (51) (**Table 1**, and **Figs. S2, S3**). A varying number of images were recorded using the EPU software in counting or super resolution mode, featuring a pixel size of 1.2 Å or 1.1 Å for C-TAC or Mor-TAC, a dose rate of 10 e/pixel/s, and an electron exposure dose of 50 e/Å^2^. Movies were captured over a duration of 8.38 seconds with the defocus range varying from -2.0 μm to -1.0 μm. Subframes of individual movies were aligned using MotionCor2 (52), While the contrast-transfer-function for each summed image was estimated using CTFFIND4 (53). From the summed images, particles were picked by blob picker, template picker and subjected to iterative 2D classification in CryoSPARC v4.2. The resulting 2D classes, exhibiting diverse orientations were further selected, manually inspected, and subjected to ab-initio reconstruction and hetero refinement. By eliminating poorly populated classes, the selected particles were then subjected with or without masked 3D classification focusing on upstream DNA region of C-TAC or Mor-TAC. Subsequently, particles from the optimal class, which displayed clear density for RNAP, DNA, C or Mor were re-processed through homogeneous refinement, non-uniform refinement, CTF-refinement, local resolution estimation, and local filtering, ultimately yielding the final map in CryoSPARC v4.2. The final particles were further subjected to particle subtraction to preserve the signal from the upstream C or Mor binding regions, followed by masked local refinements to improve the map quality and interpretability.

**Table 1.** Single particle cryo-EM data collection, processing, and model building for *E. coli* Mor-TAC and C-TAC.

| Protein-DNA complex | Mor-TAC | C-TAC |
| --- | --- | --- |
| PDB ID | 9L0X | 9L0Y |
| EMDB ID | 62727 | 62728 |
| **Data collection and processing** |  |  |
| Microscope | Titan Krios | Titan Krios |
| Voltage (kV) | 300 | 300 |
| Detector | K3 summit | K3 summit |
| Electron exposure (e/Å<sup>2</sup>) | 50 | 50 |
| Defocus range (μm) | -1.0~-2.0 | -1.0~-2.0 |
| Data collection mode | super resolution | counting |
| Physical pixel size (Å/pixel) | 1.1 | 1.2 |
| Symmetry imposed | C1 | C1 |
| Initial particle images | 412,492 | 523,383 |
| Final particle images | 99,820 | 217,572 |
| Map resolution (Å)<sup>a</sup> | 4.12 | 3.14 |
| **Refinement** |  |  |
| Map sharpening B-factor (Å) | -156 | -160 |
| Root-mean-square deviation |  |  |
| Bond length (Å) | 0.006 | 0.003 |
| Bond angle (°) | 0.723 | 0.587 |
| MolProbity statistics |  |  |
| Clashscore | 16.13 | 11.85 |
| Rotamer outliers (%) | 0.08 | 0.25 |
| Cβ outliers (%) | 0.0 | 0.0 |
| Ramachandran plot |  |  |
| Favored (%) | 94.99 | 96.92 |
| Outliers (%) | 0.00 | 0.00 |

<sup>a</sup> Gold-standard FSC 0.143 cutoff criteria.

The final mean map resolutions for C or Mor-dependent transcription activation complex, as assessed using the Gold-standard Fourier-shell-correlation method, are as follows: 3.14 Å for C-TAC, 4.12 Å for Mor-TAC, 3.97 Å for the C region in C-TAC, and 6.92 Å for the Mor region in Mor-TAC (**Table 1** and **Figs. S4, S5**).

### Cryo-EM model building and refinement

The model of RNAP and the downstream DNA, derived from the cryo-EM structure of *E.coli* RPo (PDB ID: 6OUL), as well as the predicted structure of C (ID: AF-E0J3R8-F1) or crystal structure of Mor (PDB ID: 1RR7) were fitted into the cryo-EM density map of C-TAC and Mor-TAC using Chimera (54) to generate a preliminary structural model. The model of the upstream nucleic acids was built manually using Coot (55). The coordinates were subsequently calculated and validated through real-space refinement incorporating secondary structure restraints in Coot and Phenix (v1.19.2) (56). Structures were analyzed using Chimera and PyMOL(57). The Map versus Model FSCs of the four cryo-EM maps (C-TAC, Mor-TAC) in this work were generated by Phenix. The statistics of cryo-EM refinement were summarized in **Table 1**.

### *In vitro* transcription assay

To assess the effects of Mor or C on activating transcription of the phage Mu middle gene promoter P*m* or late gene promoter P*mom*, we performed i*n vitro* transcription assays with a 96-well microplate in the transcription buffer (20 mM Tris–HCl, pH 8.0, 50 mM NaCl, 5% glycerol, 10 mM MgCl2). Each component was included in the reaction mixture (80 μl) as following: 0 or 4 μM C or C mutants (Mor or Mor mutants), 0.1 μM RNAP, 50 nM P*mom* DNA (or P*m* DNA), 0.1 mM NTP mix (ATP, UTP, GTP, and CTP), and 1 μM TO1-Biotin. Initially, RNAP, promoter DNA, and C or C mutants (Mor or Mor mutants) were incubated for 15 min at 37 ℃. Then, NTP mix and TO1-biotin were supplemented into the mixture to trigger transcription initiation for 10 min at 37 ℃. Finally, the fluorescence emission intensities were measured using a multimode plate reader (EnVision, PerkinElmer Inc) with an excitation wavelength at 510 nm and an emission wavelength at 535 nm. Relative transcription activities of the relevant derivatives were calculated using the following formula (1):

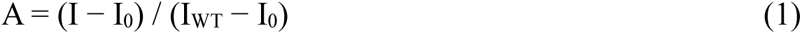

where I_WT_ and I are the fluorescence intensities in the presence of C or C mutants (Mor or Mor mutants), I0 is the fluorescence intensity in the absence of C (or Mor) protein.

### Electrophoretic mobility shift assay

Electrophoretic mobility shift assays (EMSA) of phage Mu C (or Mor) protein were performed in reaction mixtures (20 μl) containing: 30 nM P*mom* DNA (or P*m* DNA), 0 or 0.15 μM *E. coli* RNAP, and supplemented with 4 μM GST-C (or GST-Mor) in the EMSA buffer (20 mM Tris–HCl, pH8.0, 50 mM NaCl, 5 mM MgCl_2_, 5% glycerol). Firstly, the promoter DNA, *E. coli* RNAP were incubated for 10 min at 30 ℃. Then the GST-C (or GST-Mor) protein was added into the reaction mixture, and incubated for another 20 min at 30 ℃ before addition of 0.075 mg/ml heparin. Finally, the samples were applied to 5% polyacrylamide slab gels (29:1 acrylamide/bisacrylamide), electrophoresed in 90 mM Tris borate, pH 8.0, and 20 mM EDTA, and stained with 4S Red Plus Nucleic Acid Stain (Sangon Biotech, Inc.) according to the procedure of the manufacturer.

### Single-molecule FRET assay

HPLC purified DNA oligos (C-nonTem/C-Tem, and Mor-nonTem/Mor-Tem listed in **Tables S1** and **S2**, the underlined “T” is labeled with Cy3 for non-template strand and Cy5 for template strand, respectively, Sangon Biotech) were annealed at 1:1 molar ratio in the transcription buffer by heating to 95 °C for 5 min followed by slowly cooling down to room temperature in about two hours, named P*mom*-DNA and P*m*-DNA, respectively. After adding a final concentration of 0.5 mg/ml BSA, the annealed DNA constructs was aliquoted and stored at -20 °C.

In single-molecule FRET assay, the Cy3/Cy5 doubly labeled DNA substrates were tethered to the PEG surface of a flow chamber precoated with streptavidin (ThermoFisher Scientific). Experiments with DNA alone or in presence of 10 nM RNAP, or 10 nM RNAP + 100 µM NTPs, or 10 nM RNAP + 100 µM NTPs + 500 nM C protein or Mor protein, were performed in the transcription buffer containing 0.5 mg/ml BSA, 2.5 mM PCA (protocatechuic acid, Sigma-Aldrich), 50 nM PCD (protocatechuate-3,4-dioxygenase, Sigma-Aldrich) and 1 mM Trolox (Sigma-Aldrich). For the experiments with C or Mor proteins, 500 nM C or Mor proteins were incubated with surface-tethered DNA for 10 min at room temperature, respectively, followed by addition of 10 nM RNAP + 100 µM NTPs + 500 nM C protein or Mor protein in the transcription buffer containing 0.5 mg/ml BSA, 2.5 mM PCA (protocatechuic acid, Sigma-Aldrich), 50 nM PCD (protocatechuate-3,4-dioxygenase, Sigma-Aldrich) and 1 mM Trolox (Sigma-Aldrich). Data was collected at an acquisition rate of 10 Hz on a home-made TIRF-based microscope under the excitation of 532 nm laser (Coherent). Fluorescence was imaged by an objective (Apo TIRF 100x, 1.49NA, Olympus), splitted into two channels and then collected on an EMCCD (iXon Ultra 897, Andor). Raw data was extracted, and background corrected. The FRET value or the number of transcription events (RPo, RPitc or none) were analyzed manually.

## RESULTS

### Overall structures of Mor-TAC and C-TAC

To elucidate the structural basis underlying the transcription activation mechanism of the distinct Mor/C family factors, we purified *E*. *coli* RNAP, and phage Mu Mor/C proteins to high homogeneity, and prepared the phage Mu middle gene promoter P*m* scaffold and the late gene promoter P*mom* scaffold. Each of the scaffold contains a consensus –10 element, a transcription bubble, and an imperfect dyad-symmetry element (comprising A site and B site) located just upstream of the nonoptimal -35 element (**Fig. 1A** and **Fig. S4**). *In vitro* transcription assays demonstrated that Mor and C activated transcription of the phage Mu middle gene promoter P*m* and late gene promoter P*mom*, respectively (**Fig. S5A**). Electrophoretic mobility shift assays (EMSA) yielded shifted bands for Mor-TAC and C-TAC compared to the binary Mor/C-DNA complex or the RNAP-DNA complex (**Fig. S5B**). These findings are in good agreement with the previous observations that Mor recruits RNAP to P*m* rather than repositioning a prebound RNAP, a process that occurs concurrently for C-dependent repositioning of RNAP at P*mom* (39,43). Consistently, gel filtration maps and SDS-PAGE analysis of the purified phage Mu Mor-TAC and C-TAC demonstrated the proportional appearance of each protein component (**Fig. S5, C** and **D**).

**Fig. 1.**
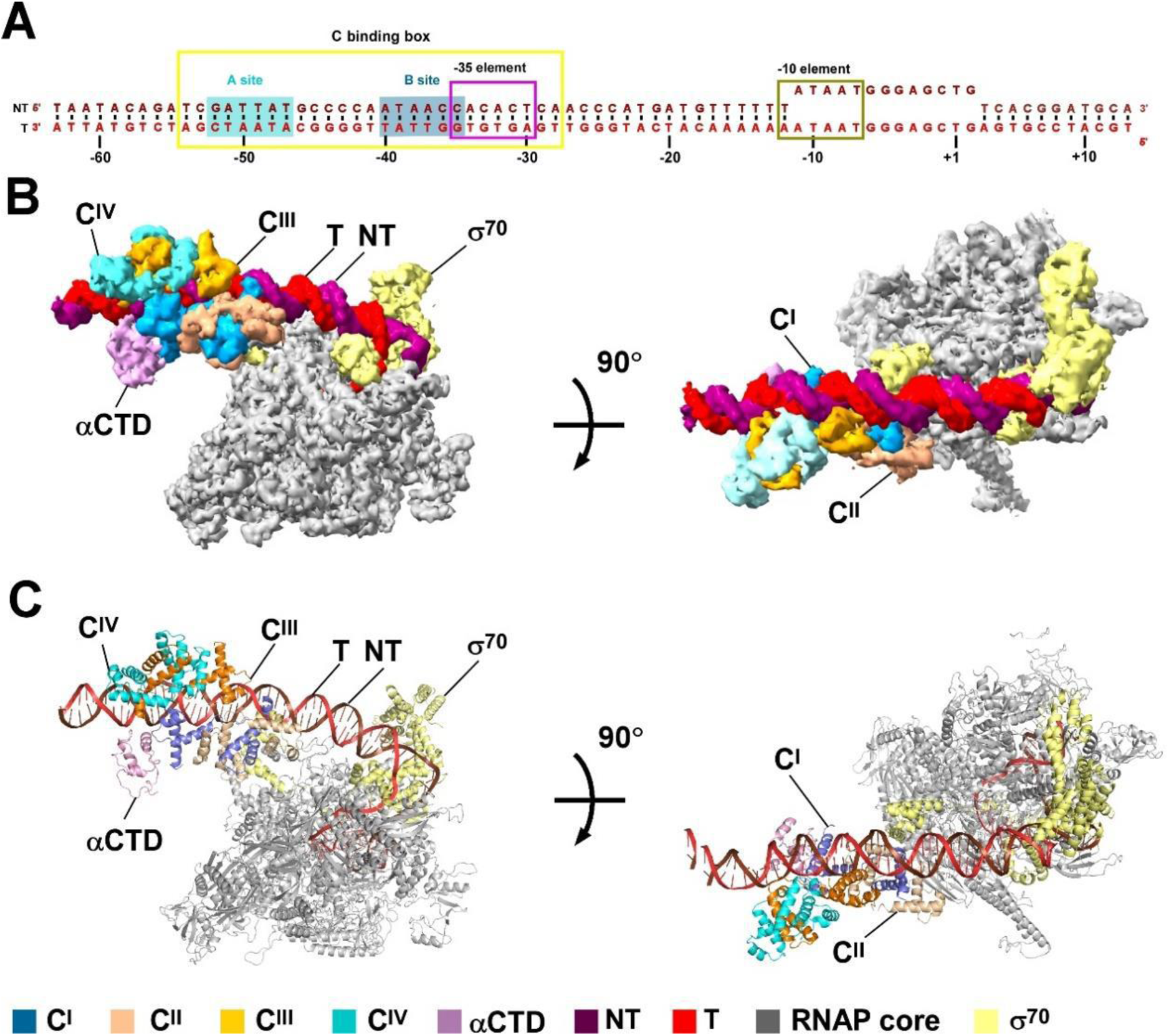
Overall structure of the phage Mu C-TAC. **(A)** DNA scaffold used in structure determination of Phage Mu C-TAC. **(B, C)** Two views of the cryo-EM density map (B) and structure model **(C)** of Phage Mu C-TAC. The EM density maps and cartoon representations of C-TAC are colored as indicated in the color key. Firebrick, NT, non-template-strand promoter DNA; red, T, template-strand promoter DNA. blue, C^I^; wheat, C^II^; orange, C^III^; Cyan, C^IV^; light blue, C binding site A; dark blue, C binding site B; yellow, *E. coli* RNAP σ^70^ subunit; violet, *E. coli* RNAP αCTD. The C binding box, -35 element, and -10 element is boxed with yellow, violet, and brown colors, respectively. The EM density maps are colored as indicated in the color key.

After data collection of the optimized TACs samples, we processed the cryo-EM data and performed model building by combination of different structural software. The cryo-EM maps of Mor-TAC and C-TAC were determined at nominal resolutions of 4.12 Å and 3.14 Å, respectively (**Fig. 1**, **Tables 1, 2**, and **Figs. S2-S4, S6, S7**). The RNAP subunits and the downstream DNA from the cryo-EM structure model of *E*. *coli* RPo (PDB ID: 6OUL) were able to be fitted into the discerned electron densities of the RNAP core regions in TACs (58). Additionally, the DDs and DBDs from the reported crystal structures of the Mor dimer (PDB ID: 1RR7) and the AlphaFold-predicted C monomer (ID: AF-E0J3R8-F1) were well fitted into the electron densities surrounding the upstream DNA in Mor-TAC and C-TAC, respectively.

**Table 2.** Single particle cryo-EM data collection, processing for focusing on Mor or C region in *E. coli* Mor-TAC and C-TAC.

| **Protein-DNA complex** | **Mor-TAC** | **C-TAC** | 721 |
| --- | --- | --- | --- |
| EMDB ID | 63410 | 63408 |  |
| **Data collection** |  |  |  |
| Voltage (kV) | 300 | 300 |  |
| Detector | K3 summit | K3 summit |  |
| Electron exposure (e/Å<sup>2</sup>) | 50 | 50 |  |
| Defocus range (μm) | -1.0~-2.0 | -1.0~-2.0 |  |
| Data collection mode | super resolution | counting |  |
| Physical pixel size (Å/pixel) | 1.1 | 1.2 |  |
| Symmetry imposed | C1 | C1 |  |
| Initial particle images | 412,492 | 523,383 |  |
| Final particle images | 99,820 | 217,572 |  |
| Map resolution (Å)<sup>a</sup> | 6.92 | 3.97 |  |

<sup>a</sup> Gold-standard FSC 0.143 cutoff criteria.

Strikingly, the Mor/C proteins bind promoter DNA as centrosymmetric tetramers in both Mor-TAC and C-TAC (**Figs. 1, 2,** and **Figs. S4, S6-S10**), which are quite different from the docked dimeric model of Mor and DNA (30,32). In contrast, these differences greatly coincide with the predicted conformation changes for transactivation of the Mor/C proteins. Though the N-terminus and C-terminus are not wholly visualized, the corresponding electron densities from each protomer intertwined together over the tetramer, suggesting positive roles of the termini in stabilizing this unique architecture (**Fig. 2**, and **Figs. S8-S10**). Moreover, there are evident overlapping densities between the Mor/C protein and the conserved domains of *E*. *coli* RNAP, indicative of extensive interactions between the Mor/C proteins and RNAP (**Fig. 3**, and **Figs. S11, S12**), which may facilitate recruitment of RNAP to the promoter region and enhance stabilization of the functional TACs. Besides, the upstream DNA in C-TAC exhibits a larger bending angle compared to that in Mor-TAC or phage λCII-TAC (59), which may provide valuable insights into C-dependent transcription activity on phage Mu late gene promoter (**Fig. 1**, and **Fig. S13**). Given the higher resolution of C-TAC, more diverse protein-protein interactions, and more complex transactivation mechanism of C compared to Mor, the structural model of C-TAC is emphasized in the following text to elucidate the distinct transcription activation mechanisms of the Mor/C family activators, using Mor-TAC as a comparison.

**Fig. 2.**
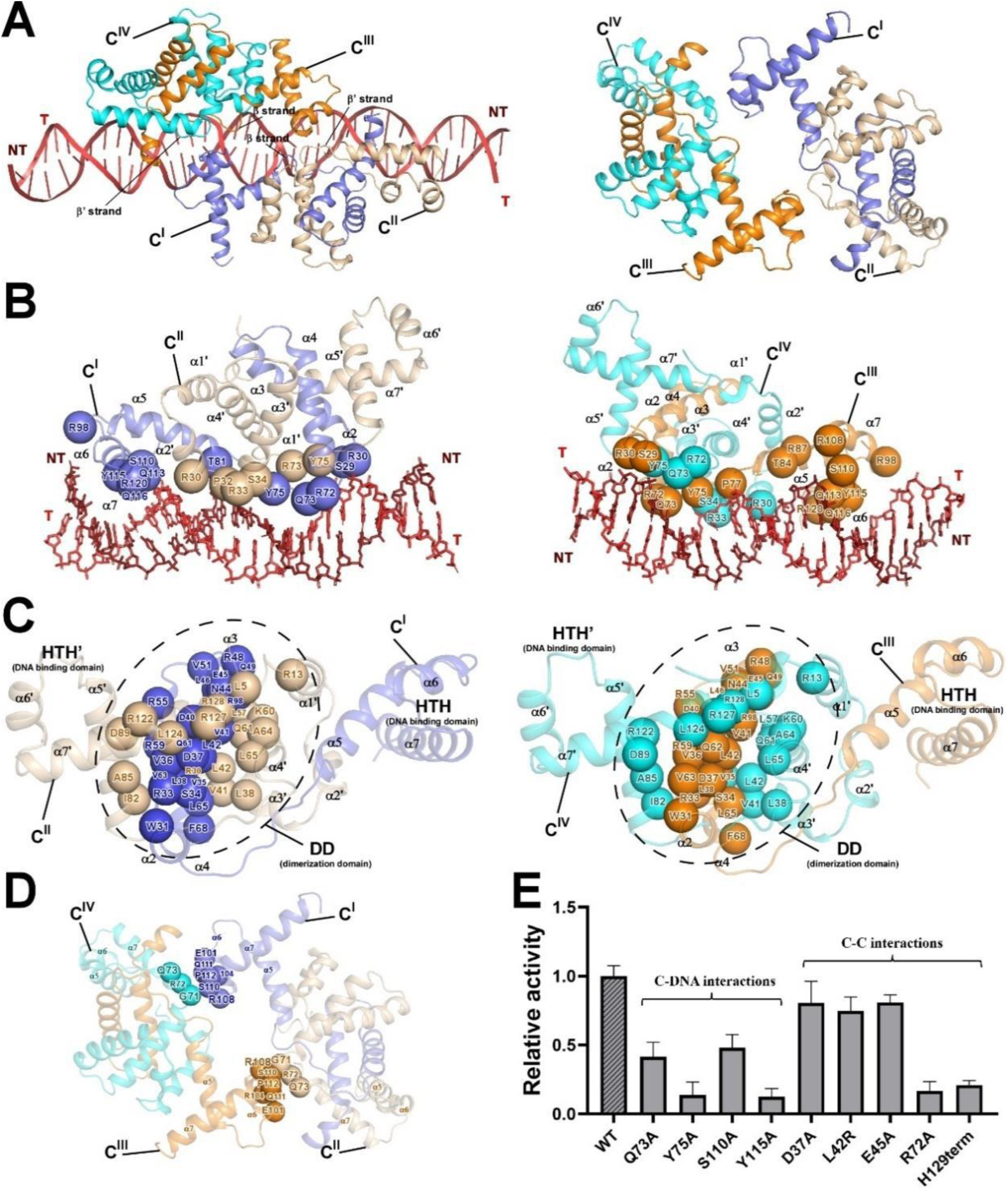
C binds promoter DNA as a centrosymmetric tetramer in the phage Mu C-TAC. **(A)** Relative locations of the centrosymmetric C tetramer at the upstream double-stranded DNA (left panel); structure of C tetramer in cartoon (right panel). **(B)** The relative locations between C^I^ and C^II^ (left panel) or C^III^ and C^IV^ (right panel) bound to the promoter DNA. The secondary structural elements involved in C tetramer are labeled, respectively. Colors of C^I^, C^II^, C^III^, C^IV^, NT, and T in A-C are shown as in Fig. 1C. **(C)** Detailed dimerization interactions between C^I^ and C^II^ (left panel) or C^III^ and C^IV^ (right panel). **(D)** Detailed tetrameric interactions between C^I^ and C^IV^ or C^II^ and C^III^. The secondary structural elements are labeled, respectively. Colors are shown as in Fig. 1. The key residues involved in B-D are shown as spheres in the corresponding colors. **(E)** Mutations of the C residues involved in DNA binding, dimeric and tetrameric interfaces display reduced transcription activities of the C protein. Data for *in vitro* transcription assays are means of 3 technical replicates. Error bars represent mean ± SEM of n = 3 experiments.

**Fig. 3.**
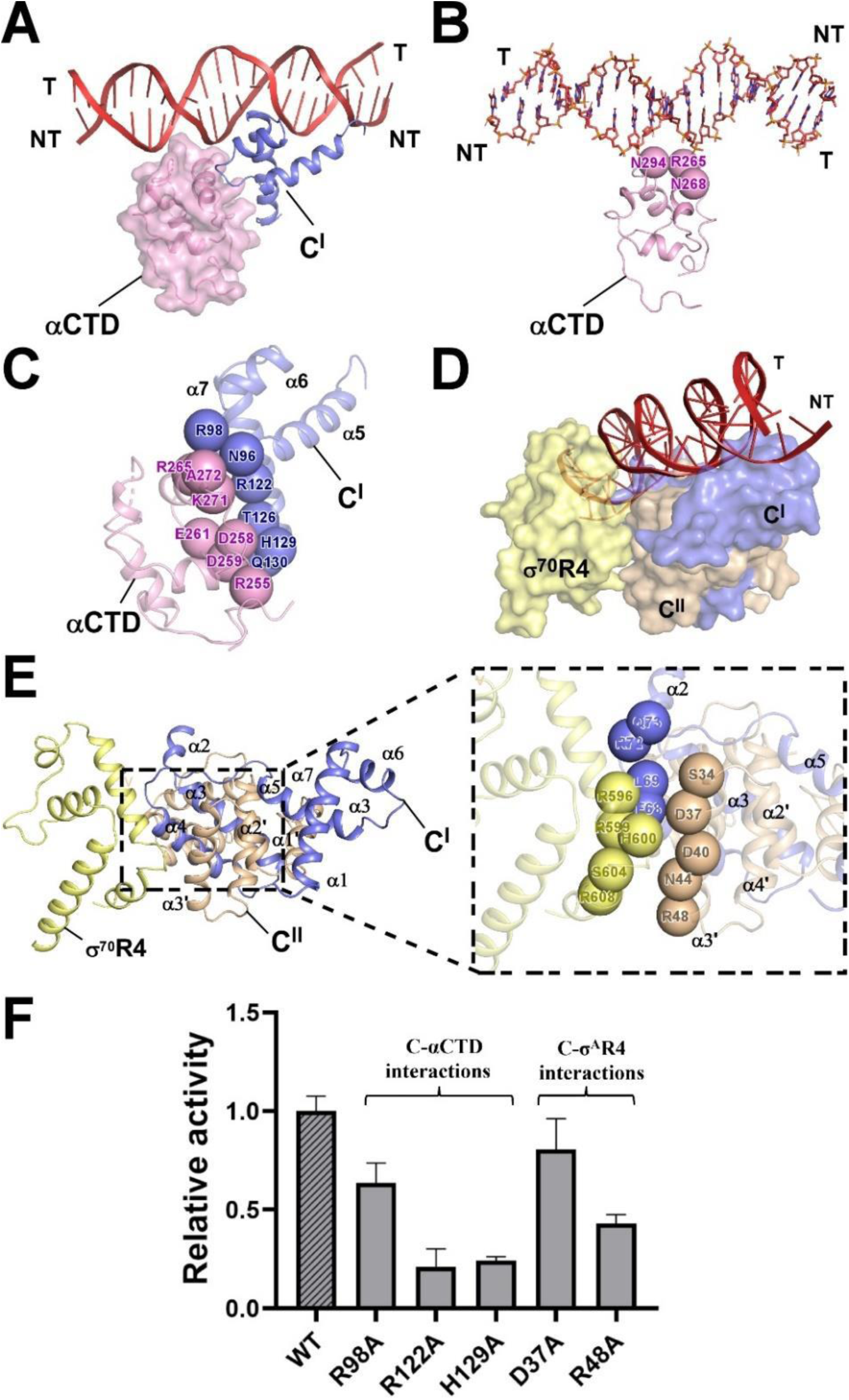
C makes extensive interactions with the conserved domains of RNAP in the phage Mu C-TAC. **(A)** Relative locations of *E. coli* RNAP αCTD, C^I^, and the upstream double-stranded DNA. *E. coli* RNAP αCTD, C^I^ are also represented as magenta or orange cartoon, respectively. RNAP αCTD is also shown in surface. **(B)** Detailed interactions between *E. coli* RNAP αCTD and the upstream double-stranded DNA, with key residues involved in αCTD shown as magenta spheres. **(C)** Detailed interactions between *E. coli* RNAP αCTD and C^I^, with key residues involved in αCTD and C^I^ shown as magenta and blue spheres, respectively. **(D)** Relative locations of *E. coli* RNAP σ^70^R4, C^I^, C^II^, and the upstream double-stranded DNA. C^I^, C^II^ and RNAP σ^A^R4 are shown in surface and colored as in **Fig.1C**. **(E)** Critical interactions between the C^I^-C^II^ dimer and *E. coli* RNAP σ^70^R4. Residues involved in the magnified interfaces (right panel) between the C^I^-C^II^ dimer and RNAP σ^A^R4 are shown as yellow (RNAP σ^A^R4), blue (C^I^), and wheat (C^II^) spheres, respectively. **(F)** Mutation of residues of C involved in the above interfaces suppressed *in vitro* transcription activity. Data for *in vitro* transcription assays are means of three technical replicates. Error bars represent ± SEM of n = 3 experiments.

### Interactions mediate DNA engagement and tetramerization in Mor-TAC and C-TAC

In Mor-TAC and C-TAC, both the Mor and C centrosymmetric tetramers undergo significant conformational changes to facilitate DNA engagement and tetramerization through newly uncovered interactions (**Fig. 2**, and **Figs. S2, S6-S10**), which align well with predictions from earlier biochemical and crystal structural studies (30,32).

In C-TAC, the downstream C dimer, intertwined by C^I^ and C^II^, and the upstream dimer, associated with C^III^ and C^IV^, coordinately assemble into a centrosymmetric tetramer that extends to occupy nearly three helical turns upstream the potential -35 element (**Fig. 2A**). This positioning coincides with the far apart DNA binding sites (26 bp) observed in previous footprinting assays (30,36). In contrast to the docked Mor-DNA structural model, both the conserved C-terminal HTH motif (comprising residues R98, S110, Q113, Y115, Q116, and R120) of C^I^ constituted by helices α5-α7, and the anti-paralleled β-strands (including residues R72, R73, Y75, P77, and T81) from C^III^ and C^IV^ move away from the dimerization domain (DD) and insert into the upstream DNA major groove from opposite directions (**Fig. 2B**). Additionally, two flanking loops connecting helix α2 and helix α3, consisting of residues S29, R30, P32, R33, and S34, make contacts with the DNA backbones of the adjacent DNA minor grooves. By analogy, the centrosymmetric anti-paralleled β-strands from C^I^ and C^II^ and the conserved C-terminal HTH motif of C^III^ specifically interact the downstream DNA major groove, accompanying by two flanking loops interacting with the adjacent DNA minor grooves (**Fig. 2B**). These interactions allow for stable DNA engagement of the Mor/C proteins at the corresponding binding box, which are further favored by the centrosymmetric tetramer architecture. Except for the clustered hydrophobic and polar interactions involved in each DD of the C dimers (**Fig. 2C**), which exhibit similarities with those observed in the crystal structure of the Mor dimer, polar contacts between helix α6 of C^I^ and the β-strand of C^IV^, as well as between helix α6 of C^III^ and the β-strand of C^II^ contribute to the tetramerization of the C^I^-C^II^ and C^III^-C^IV^ dimers (**Fig. 2D**), thereby enhancing formation and stability of the centrosymmetric tetramers within the TACs. Notably, mutations of the C residues that are critical for DNA binding and for dimeric and tetrameric interactions compromise the *in vitro* transcription activities of the C protein (**Fig. 2E**). Specifically, mutations at residues Y75, Y115, R72, and truncation of the C-terminus (H129term) nearly abolished the relative transcription activity of C, underscoring the essential roles of these residues and the previously uncharacterized C-terminus in sustaining C-dependent transcription activation (30).

Similarly, comparable protein-DNA and protein-protein interactions are observed in the structure of Mor-TAC. Mutations affecting these interfaces, as well as truncation of the acid N-terminus (del_N10) or the basic C-terminus (L119term), result in defects in the transcription activities of the Mor protein (**Figs. S9, S10**), thereby highlighting the significance of these interfaces in promoting Mor-dependent transcription activation.

### Interactions mediate RNAP recruitment in Mor-TAC and C-TAC

In addition to the EMSA results which displayed apparent roles in recruiting RNAP (**Fig. S5B**), structural analysis of both Mor-TAC and C-TAC also provide detailed evidences for RNAP recruitment of the Mor/C proteins (**Fig. 3** and **Figs. S11**, **S12**).

Biochemical and genetic studies have demonstrated that both the middle and late gene promoters of Mu phage contain UP elements (AT-rich element) that may be specifically recognized by the conserved domain of RNAP αCTD (38,40). Mutagenesis experiments also indicate that σ^70^R4 is required for Mor/C-dependent transcriptional activation (25,42). Supporting these findings, in C-TAC, the conserved domain αCTD of RNAP not only interacts with the UP elements of promoter DNA through its 265 determinant (including residues R265, N268, and N294), but also establishes polar contacts with the C-terminal of helix α7 and the loop connecting helices α5 and α6 from C^I^, respectively (**Fig. 3, A-C**). Besides, the residues from the β-strand of C^I^ and helix α3’ of C^II^ interact with the two conserved C-terminal helices of σ^70^R4 (**Fig. 3, D** and **E**), exhibiting a buried surface area of approximately 125 Å^2^. Accordingly, substitutions of these residues involved suppressed *in vitro* transcription activity of the C protein, particularly for residues R48, R122, and H129 (**Fig. 3F**). Likewise, structural analysis and mutagenesis of Mor-TAC yielded comparable results (**Fig. S11**). Notably, the residue C67 from the C^I^-C^II^ dimer is likely to make contacts with the residues R599 and I905 from the conserved β flap tip helix (βFTH) of RNAP, an interface not observed in Mor-TAC (**Fig. S12**). These extensive protein-protein interactions involving C may enhance its higher capability to recruit RNAP, thereby facilitating the formation of a more stable C-TAC complex compared to Mor-TAC.

### Single-molecule FRET assays show C significantly promotes RPitc formation

To further assess whether C activates transcription initiation in subsequent processes following the formation of TAC as previously proposed, we conducted dynamic single-molecule Förster resonance energy transfer (smFRET) assays by using Mor as a control. We prepared doubly labeled DNA substrates (Cy3 at -4 position of the non-template strand and Cy5 at +17 position of the template strand, **Fig. 4A**). The FRET between these two dyes was utilized to characterize the changes in their distances during interacting with RNAP, which enable us to monitor the transition states of RNAP during transcription initiation in the presence of either the C protein or the Mor protein (60).

**Fig. 4.**
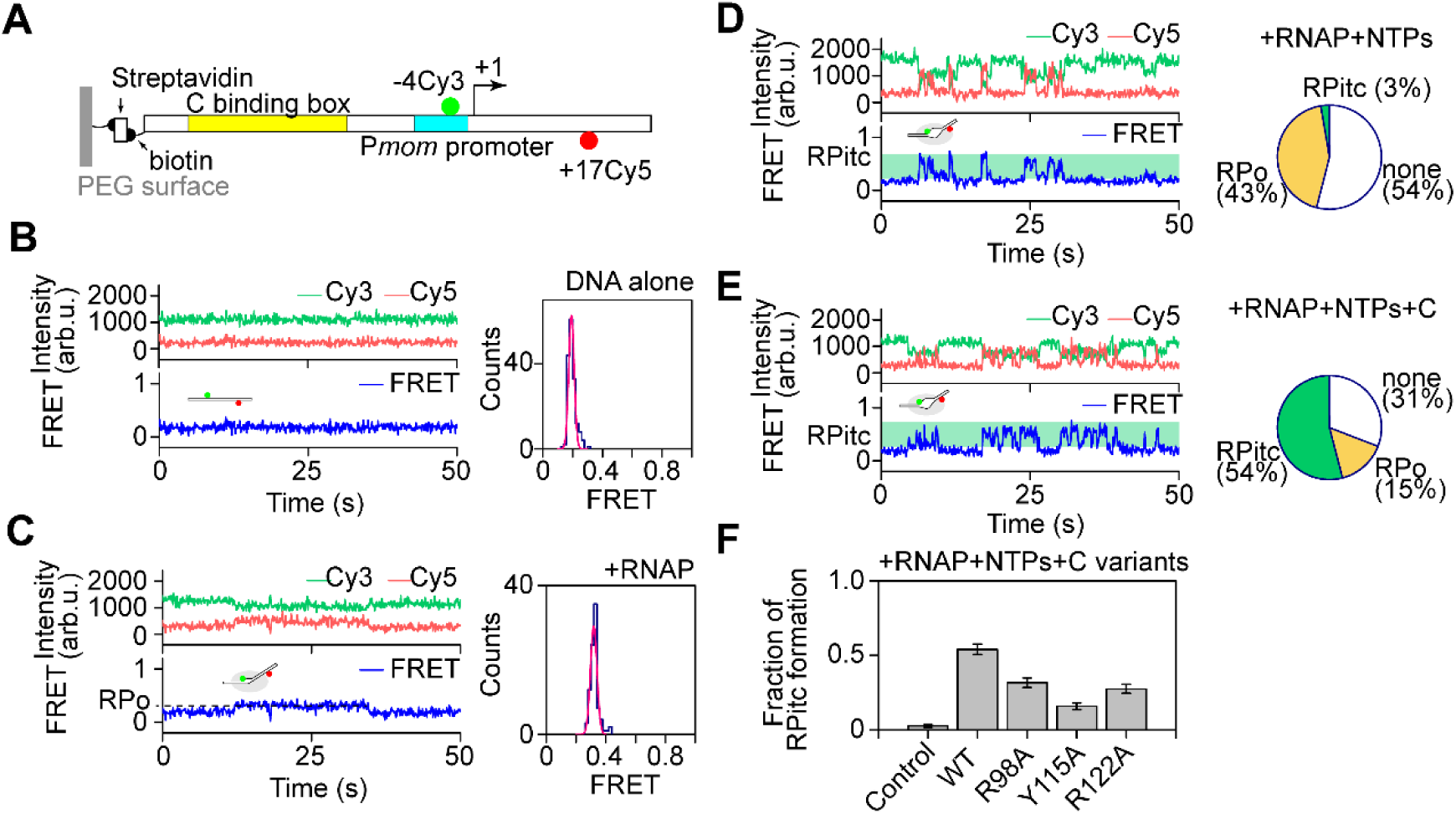
Characterization of C protein on RNAP transcription activity via single-molecule FRET assay. **(A)** Schematic of the DNA substrate for the single-molecule FRET assay. Typical trajectories representing the intensities of Cy3 and Cy5 and thus the FRET for each reaction condition (left panels): Surface-tethered DNA alone **(B),** Surface-tethered DNA + 10 nM RNAP **(C)**, Surface-tethered DNA + 10 nM RNAP + 100 µM NTPs **(D)**, and Surface-tethered DNA + 10 nM RNAP + 100 µM NTPs + 500 nM C protein **(E)**, respectively. FRET distributions of each condition are fit to a single-Gaussian function yielding peaks at 0.19 ± 0.002 (SEM, N = 166, right panel, B), and 0.32 ± 0.002 (SEM, N = 89, right panel, C). Pie charts representing the fractions of DNA molecules occurred in RPo (N = 114 for D, and 67 for E, yellow), RPitc (the initial transcription complex, N = 7 for D, and 240 for E, green), and without RPo or RPitc complexes (N = 142 for D, and 136 for E, white) in the absence and the presence of C protein, respectively, (right panels). **(F)** Mutation of the C residues involved in C-DNA, C-αCTD interfaces suppress fractions of RPitc formation.

As shown in **Fig. 4**, surface-tethered DNA molecules alone yield a mean FRET value of 0.19 (**Fig. 4B**). This value increases to 0.32 upon the addition of RNAP (**Fig. 4C**), indicating a distance reduction between the Cy3/Cy5 dyes caused by bending and unwinding of promoter DNA, resulting in the formation of the RPo complex. When RNAP and NTPs are added, 121 out of 263 DNA molecules remain in the RPo state, however, 7 out of these 121 molecules exhibit abrupt FRET increases that exceed those observed in the RPo state, suggesting that RNAP transitions into the initial transcription complex (RPitc) state (**Fig. 4D**). To investigate the influence of C on the transcription activity of RNAP, C was incubated with the DNA substrate on a PEG surface at room temperature for 10 min, followed by the addition of RNAP, NTPs and C. As anticipated, 307 out of 444 molecules formed the RPo state. Notably, 240 of these 307 molecules transitioned into RPitc states, indicating a significant role of C in promoting transcription initiation (**Fig. 4E**). Consistently, mutations of the relevant C residues (R98A, Y115A, and R122A) resulted in a decreased fraction of RPitc formation (**Fig. 4F**), indicative of the importance of these residues in activating transcription during the later process. In contrast, similar experiments conducted with the Mor protein on the phage Mu middle gene promoter demonstrated a weaker effect of Mor in facilitating RNAP transcription initiation compared to C (**Fig. S14**).

Collectively, these data suggest that C function as a transcriptional activator in a manner different from Mor during the transcriptional initiation phase after TAC formation, which further supports the previous observations from a dynamic perspective that C also facilitates transcriptional activation by enhancing promoter clearance (43,44).

## DISCUSSION

Recently, the emerging cryo-EM single particle reconstruction technology has significantly accelerated structural investigation of phage transcriptional regulation, including large intermediate complexes difficult to determine by X-ray crystallography or NMR (1). Here, by utilizing cryo-EM to investigate the model system phage Mu, we have elucidated long-standing questions regarding the transcription activation mechanisms of the Mor/C family activators through the following new findings.

First, the Mor/C family transcription activators form distinct centrosymmetric tetramers that elegantly stabilize Mor-TAC and C-TAC via extensive protein-DNA and protein-protein interactions (**Figs. 1, 2** and **Figs. S2, S6-S10**). Unlike most transcription factors that utilize HTH motifs to engage with the major and minor grooves of DNA, two Mor/C dimers simultaneously interact with two adjacent major grooves concurrently through both their conserved HTH motifs and the unique anti β-strands, accompanying by contacts made through short helices α2 with two bilateral minor grooves. Such protein-DNA interactions are also favored by substantial interactions between Mor/C and the conserved domains (αCTD, σ^70^R4, and β FTH) of RNAP (**Figs. 3, 5A** and **Figs. S11, S12**). Collectively, these interactions lead to significant bending of the upstream promoter DNA in TACs, particularly in C-TAC, compared to the canonical RNA polymerase open complex (RPo) (**Fig. 5**). These findings, consistent with previous biochemical and genetic assays, provide new detailed evidences for the ’recruitment mechanism’ of Mor-dependent transcription activation of phage Mu middle genes, as well as the dual ’recruitment-repositioning mechanisms’ for C-dependent transcription activation of phage Mu late genes.

**Fig. 5.**
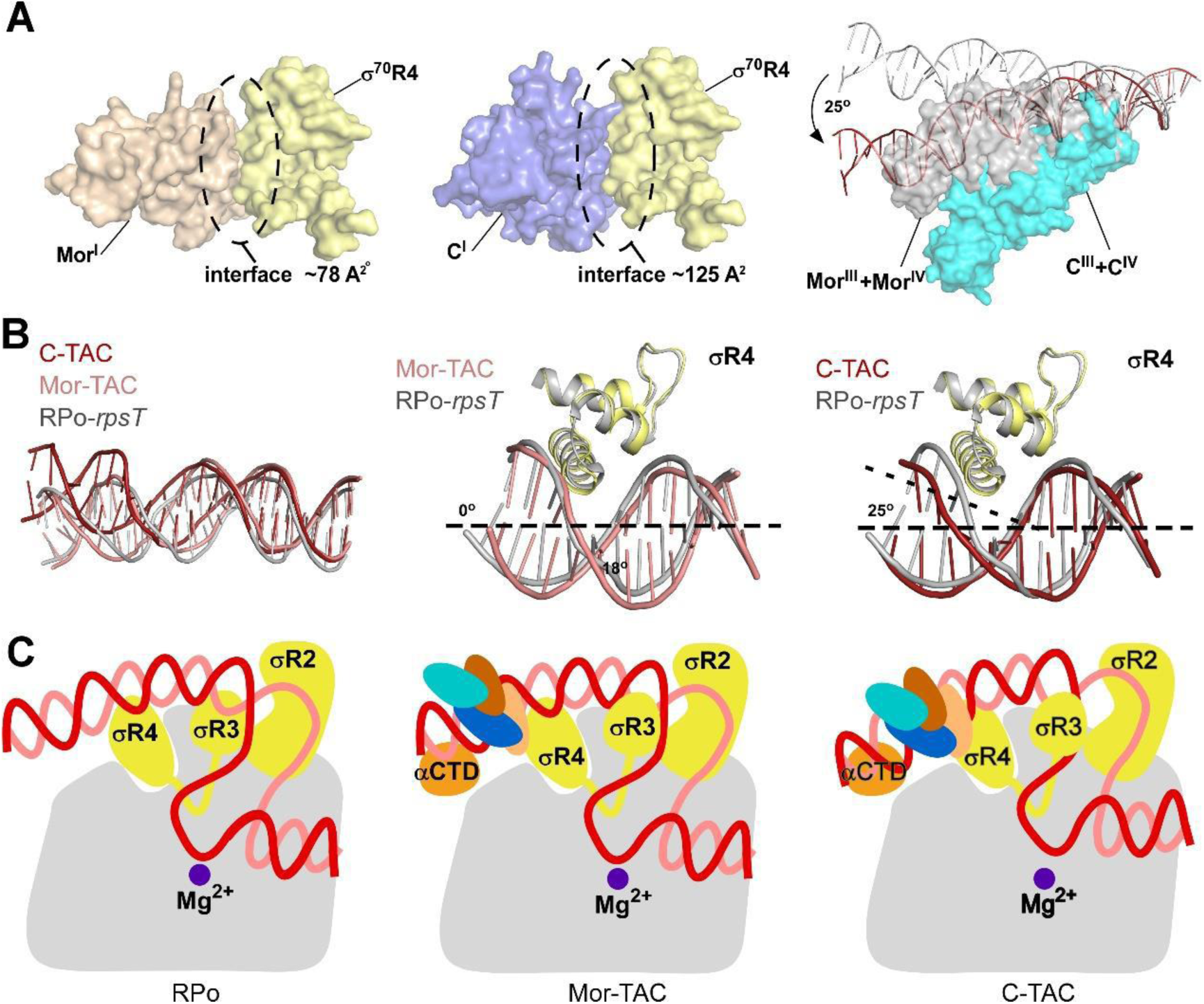
Proposed transcription activation model for the phage Mu Mor/C family proteins. **(A)** The interface of σ^70^R4 with Mor (left panel) or C (middle panel), Mor^I^ or C^I^ and σ^70^R4 were represented as surface in wheat, blue and yellow; alignment between the cryo-EM structures of C-TAC and Mor-TAC on the upstream DNA (right panel), the surface representations of Mor^III^ and Mor^IV^ are colored as in gray. The surface representations of C^III^ and C^IV^ are colored as in cyan, the upstream DNA of Mor-TAC or C-TAC was shown as cartoon in gray or red, respectively; **(B)** Alignment between the cryo-EM structures of RPo-*rpsT* (PDB ID: 6OUL, gray) and Mor-TAC(salmon) or C-TAC(firebrick) on the upstream DNA (left panel), the structure of Mor-TAC, C-TAC was superimposed on the structure of RPo-*rpsT*, The superimpositions were performed using σ^70^R4 atoms. (**C)** Proposed transcription activation model for the phage Mu Mor/C family proteins. Compared to a canonical RPo (left panel), central symmetrically arranged Mor homotetramer hijacks *E. coli* RNAP through interacting with its conserved domains (αCTD and σ^70^R4) and recruits RNAP to activate transcription of the phage Mu middle promoter through engaging promoter DNA (middle panel). In contrast, central symmetrically arranged C homotetramer makes diverse interactions with the conserved domains (αCTD, σ^70^R4 and β FTH) of RNAP and significantly bends the upstream promoter DNA, which cooperatively recruit RNAP to activate transcription of the phage Mu late promoter (right panel).

Second, significant conformational changes are observed in the homologous Mor-TAC and C-TAC as compared to the crystal structure of the Mor dimer and the docked binary model of Mor and DNA. These changes allow for large occupancy of the Mor/C family activators on promoter DNA, as proposed by biochemical assays (30–32,34,36,44). Unlike other monomeric and dimeric transcription factors, the tetrameric architecture of both Mor and C offers unique advantages: the downstream dimer is essential for interaction with the promoter B site and the conserved domains of RNAP, while the upstream dimer bound to the promoter A site may obstruct additional binding from other transcriptional activators or repressors, thereby significantly enhancing the efficiency of transcription initiation. The centrosymmetric tetramer arrangement differs from the reported tandem GlnR tetramer observed in *Mycobacterium tuberculosis* GlnR-TAC (61), presenting a novel activation mechanism for transcription factors. Third, the structural, biochemical and kinetic single-molecule FRET assays provide new insights into the distinctions between Mor-TAC and C-TAC. Due to establishing more abundant interactions with the conserved domains (σ^70^R4 and β FTH) of RNAP, C significantly alters the DNA curvature in the upstream region of C-TAC compared to Mor (**Fig. 5A**). This alteration is indicated as a prerequisite for σ^70^R4 binding in the other reported TACs, as well (18,49,59). Though σ^70^R4 and the suboptimal -35 element of Mor-TAC superimpose well on the structure of RPo-*rpsT* (PDB ID: 6OUL), the greater DNA bending angle in C-TAC, resulting from the binding of C, extends the distance between σ^70^R4 and the DNA (**Fig. 5B**), thereby weakening the reciprocal interactions. This may facilitate subsequent RPitc formation, promoter clearance, and σ escape, which were kinetically verified by single-molecule FRET assays (**Fig. 4** and **Fig. S14**). In contrast to a canonical RPo or TAC with dimeric transcription activators, Mor and C activate transcription of the phage Mu middle and late genes by acting as distinctive centrosymmetric tetramers (**Fig. 5C**).

In contrast to the structure of phage λCII-TAC, this work highlights different intriguing characteristics of the Mor/C family activator-dependent transcription activation: (i) These activators adopt as unique centrosymmetric tetramers and maintain stability of the TACs through more viable and diverse interfaces with both DNA and the conserved domains (αCTD, σ^70^R4, and β FTH) of RNAP than λCII (**Fig. S13**). Regarding the DNA binding characteristics as reviewed recently, the Mor/C family activators should be classified as class I transcription factors which bind upstream of promoter -35 element, while the λCII is categorized as a class IV transcription factor that binds both upstream and downstream of promoter -35 element. (ii) The different DNA bending angles observed in Mor-TAC and C-TAC may provide new structural insights into the conformational changes that occur during transcription initiation, reflecting the complexity of transcription regulation in the middle and late genes of phage Mu. (iii) By combing the single molecule FRET assay with the biochemical and structural observations, this study enhances our understanding on the molecular transcription mechanisms of the Mor/C family activators, which elegantly regulate the phage Mu lytic cycle by improving the specificity and efficiency of transcription activation.

In summary, our findings reveal the unique transcription activation mechanism of the tetrameric Mor/C family activators, unraveling a novel mode of phage hijacking and bacterial transcription regulation. The key regulatory elements involved in both our and the previous phage TACs may offer new targets to effectively control phage gene expression for specific phage therapy.

## DATA AVAILABILITY

The cryo-EM maps and model coordinates have been deposited under the accession numbers: EMDB: 62727 and PDB: 9L0X for the phage Mu Mor-TAC, EMDB: 62728 and PDB: 9L0Y for the phage Mu C-TAC. All data are available in the main text or the supplementary materials.

## SUPPLEMENTARY DATA

The online version contains supplementary material available at XXX.

## AUTHOR CONTRIBUTIONS

J. S., W. L., and S. W. conceived the project. Z.H. Y., Y.R. H., L.X. X., S.M. X., L.H. X., W. C., Z.Z. F., Q. S., and J. S. prepared the protein samples, performed the biochemical assays, single-molecule FERT assays, and cryo-EM sample assembly. L.Q. X., J. S., and S.M. X prepared cryo-EM samples and data acquisition. W. L. and F. Y. analyzed the cryo-EM data and determined the structures. All authors contributed to data analysis. J. S., W. L., W. C., and S. W. wrote the paper with input from all coauthors.

## ACKNOWLEDGEMENTS

We appreciate Shenghai Chang at the Center of Cryo-Electron Microscopy in Zhejiang University School of Medicine, Guangyi Li, Liangliang Kong, Jialin Duan, and Yun Song of the Electron Microscopy System at the National Facility for Protein Science in Shanghai (NFPS), Shanghai Advanced Research Institute, Chinese Academy of Sciences, China for providing technical support and assistance in data collection. We thank the Experiment Center for Science and Technology, Nanjing University of Chinese Medicine for experimental assistance. We thank the Core Facilities, Zhejiang University School of Medicine for technical support.

